# Prolific Induction of IL-6 in Human Cells by SARS-CoV-2-derived Peptide is Attenuated by Recombinant Human Anti-inflammatory Cytokines made *in planta*

**DOI:** 10.1101/2021.09.14.460246

**Authors:** Pieter H. Anborgh, Igor Kolotilin, Nisha Owens, Abdulla Azzam Mahboob

## Abstract

Development of efficient therapies for COVID-19 is the focus of intense research. The cytokine release syndrome was underlined as a culprit for severe outcomes in COVID-19 patients. Interleukin-6 (IL-6) plays a crucial role in human immune responses and elevated IL-6 plasma levels have been associated with the exacerbated COVID-19 pathology. Since non-structural protein 10 (NSP10) of SARS-CoV-2 has been implicated in the induction of IL-6, we designed Peptide (P)1, containing sequences corresponding to amino acids 68-96 of NSP10, and examined its effect on cultured human cells. Treatment with P1 strongly increased IL-6 secretion by the lung cancer cell line NCI-H1792 and the breast cancer cell line MDA-MB-231 and revealed profound cytotoxic activity on Caco-2 colorectal adenocarcinoma cells. Treatment with P2, harbouring a mutation in the zinc knuckle motif of NSP10, caused no IL-6 induction and no cytotoxicity. Pre-treatment with plant-produced human anti-inflammatory cytokines IL-37b and IL-38 effectively mitigated the induction of IL-6 secretion. Our results suggest a role for the zinc knuckle motif of NSP10 in the onset of increased IL-6 plasma levels of COVID-19 patients and for IL-37b and IL-38 as therapeutics aimed at attenuating the cytokine release syndrome.

## INTRODUCTION

The Severe Acute Respiratory Syndrome Coronavirus 2 (SARS-CoV-2) is a highly transmissible pathogenic coronavirus. It is responsible for the coronavirus disease 2019 (COVID-19) pandemic, resulting in threats to human health worldwide(WHO, 2020). The pathogenesis of the SARS-CoV-2 in humans varies in manifestation from mild cold-like symptoms to severe respiratory failure(Hu et al., 2020). Upon binding to epithelial cells in the respiratory tract, the virus starts to replicate and eventually migrates down to the alveolar epithelial cells in the lungs, where its replication increases significantly, inducing strong immune responses(Pedersen and Ho, 2020),(Tang et al., 2020). This increase in replication is accompanied by the “cytokine storm”, which brings about respiratory distress syndrome and respiratory failure. The resulting over-exuberant immune response is considered the main cause of death in COVID-19 patients (Tian et al., 2020). With the newly emerging variants of the SARS-CoV-2 virus displaying resistance towards neutralizing antibodies, hence threatening the efficacy of vaccines(Liu et al., 2020b), alternative therapeutic approaches are more needed than ever.

The likelihood of a more severe manifestation of the viral infection in COVID-19 patients appears to correlate with the plasma levels of certain cytokines(Pedersen and Ho, 2020). Of a particular interest is the pleiotropic cytokine Interleukin-6 (IL-6) and its role in SARS-CoV-2 infections, and viral infections in general(Velazquez-Salinas et al., 2019). It has been well-established that the levels of IL-6 increase during the acute phase of infection with vesicular stomatitis virus (VSV) and this increase was associated with higher virulence in pigs(Velazquez-Salinas et al., 2018). Hospitalized patients infected with Andes virus (ANDV) displayed significantly elevated levels of IL-6, and the magnitude of the increase correlated with the severity of symptoms(Angulo et al., 2017). *In vitro* studies in cells revealed that a recombinant Spike (S) protein of SARS-CoV strongly induced production of IL-6 in murine macrophages(Wang et al., 2007). IL-6 knockout mice showed high mortality when challenged with sub-lethal doses of H1N1 influenza virus, while WT mice recovered from the infection(Dienz et al., 2012). Importantly, the serum levels of IL-6 are markedly increased during the SARS-CoV-2 infection and IL-6 was proposed to be a reliable predictive biomarker of the severity of the COVID-19 in hospitalized patients(Pedersen and Ho, 2020),(Conti et al., 2020a),(Chen et al., 2020; Liu et al., 2020a). Produced by various cell types, IL-6 induces a signaling cascade involving the JAK/STAT3 pathway to regulate transcription of a multitude of genes involved in cellular signaling and regulation of gene expression(Mauer et al., 2015; Wang et al., 2013). With its crucial involvement in both pro- and anti-inflammatory processes, IL-6 was attributed a central role in the regulation of immune responses(Scheller et al., 2011).

Certain proteins of the SARS-CoV-2 virus have been implicated in inducing the increase of IL-6 including the S protein, Nucleocapsid (N) protein, and Non-Structural Protein 10 (NSP10)(Gordon et al., 2020). The effects that some of these proteins exerted on IL-6 have been known since their homologous counterparts were discovered in SARS-CoV(Wang et al., 2007). The NSP10 protein in coronaviruses appears to constitute a cofactor in the methyltransferase complexes it forms with NSP14 and NSP16(Wang et al., 2015). The crystal structure of the NSP10/NSP16 complex of SARS-CoV-2 was solved in a recent study(Krafcikova et al., 2020). Previously, peptides derived from the NSP10 sequence involved in the interaction with NSP16 of Murine Hepatitis Virus (MHV) were reported to be successful inhibitors of NSP16 2’-O-Methyltransferase activity *in vitro*, as well as rescuing mice infected with MHV(Wang et al., 2015). Peptides derived from the region of NSP10 that interacts with NSP16 of SARS-CoV virus were demonstrated to inhibit the 2’-O-methyltransferase activity in cell-free biochemical assays(Ke et al., 2012). While the potential of NSP10-derived peptides of SARS-CoV to inhibit 2’-O-methyltransferase has not been tested in cell culture, the full length NSP10 protein is known to induce increases in IL-6 levels (Gordon et al., 2020). Interestingly, the region of the NSP10 protein that forms the interaction surface with NSP16 contains a zinc “knuckle” motif. Such motifs are known to correlate with activation of IL-6. In osteoarthiritic mice, the suppression of the ZCCHC6 protein containing this domain correlated with a reduction in IL-6 expression(Ansari et al., 2019). On the other hand, expression of ZCCHC6 in osteoarthritis chondrocytes correlated with an increase in IL-6(Akhtar et al., 2014).

Elucidating the molecular mechanisms underlining SARS-CoV-2-induced pathology and searching for potential efficacious treatments to COVID-19 represent the urgent focus of the worldwide scientific community. The regulation of the bodily inflammatory responses is exerted by an intricate network of various types of mediator molecules produced by and exchanged between different cells of the immune system (Cronkite and Strutt, 2018). The IL-1 cytokine superfamily plays a crucial role in immune system homeostasis, various autoimmune pathologies, and autoinflammation(Mantovani et al., 2019). Two relatively recently discovered members of the IL-1 cytokine superfamily, IL-37 and IL-38, exhibit profound anti-inflammatory activities(Nold et al., 2010),(Lin et al., 2001). A plethora of scientific studies aimed at elucidating the biological roles of these cytokines demonstrated their pivotal action in both innate and adaptive immune responses, anti-tumor activity, and their essential involvement in mechanisms underlying diverse pathological conditions and autoimmune disorders, thus positioning those two cytokines as promising candidates for development as prospective therapeutic agents(Mei and Liu, 2019),(Allam et al., 2020),(Yang et al., 2019),(Ummarino, 2017),(Xu and Huang, 2018),(Xie et al., 2019). The use of both IL-37 and IL-38 has been proposed as a valid therapeutic approach looking to mitigate SARS-CoV-2 infection-associated immunopathology and control the acute detrimental pulmonary inflammation seen in COVID-19(Conti et al., 2020b). Recent clinical findings linked the elevated plasma levels of IL-37 as an early response in SARS-CoV-2–infected patients with a positive clinical prognosis and earlier hospital discharge, whereas lower IL-37 early responses predicted severe illness. Higher blood IL-37 levels in those patients correlated with reduced IL-6 and IL-8 levels. Furthermore, ACE2-transgenic mice infected with SARS-CoV-2 showed alleviation of lung tissue damage, when treated with recombinant IL-37(Li et al., 2020a). Importantly, the use of Tocilizumab (a neutralizing antibody raised against human IL-6) has been reported to be safe in human trials, albeit with mixed results of efficacy, suggesting exploration of other therapeutic routes for IL-6 mitigation(Masiá et al., 2020).

In the current study we employed two peptides, P1 and P2, derived from the SARS-CoV-2’s NSP10 protein region that forms the interaction surface with NSP16. The sequence of P1 was completely homologous to NSP10’s amino acids 68-96, while P2 contained the amino acid substitution, Histidine80 to Arginine (H80R), which was designed to disrupt the zinc knuckle motif. Both peptides were engineered with an N-terminal 14 amino acid sequence corresponding to the protein transduction domain of the HIV’s Trans-Activator of Transcription (TAT) protein to allow penetration of the cell membrane. Based on previous studies demonstrating a role of full length NSP10 in stimulating the secretion of IL-6 lung epithelial A549 cells(Li et al., 2020b), we hypothesized that treating cultured human cells with the designed peptides P1 and P2 could help elucidate the involvement of the zinc knuckle motif of the viral NSP10 in increased IL-6 secretion. Interestingly, upon application of the peptides onto human lung cells a profound induction of IL-6 secretion was observed in the case of P1, but not P2. We further hypothesized that the increases in IL-6 secretion could be mitigated by the use of recombinant antiinflammatory cytokines, such as IL-37b and IL-38. We found that treatment with recombinant IL-37b and IL-38, produced in engineered tobacco plants, significantly reduced the levels of IL-6 induced by P1 in human lung cells, demonstrating their potential application as treatment against the cytokine storm caused by COVID-19.

## RESULTS

### Design of NSP10-derived peptides

We designed two peptides (P1 and P2) derived from NSP10 from SARS-CoV-2. P1 contains the CCHC Zinc finger motif of NSP10. As a control, P2 differed from P1 in only one amino acid at position 26 (Table 1), replacing the histidine of the CCHC motif with an arginine (corresponding to substitution H80R in full length NSP10). Multi-Conformer Continuum Electrostatics (MCCE) calculations suggested that the binding of zinc to the coordination site is greatly affected (by over 90%) due to this mutation (Supplemental Figure 1). In keeping with previous work, both peptides were conjugated to a 14 amino acid long HIV TAT sequence to allow for cell membrane penetration. Figure 1 shows the alignment of the sequences of NSP10 from SARS-CoV, SARS-CoV-2, MHV and Middle East Respiratory Syndrome (MERS)-CoV. NSP10 sequences of SARS-CoV and SARS-CoV-2 are identical with respect to the region of interest, while sequences from MHV and MERS-CoV share a Proline to Valine substitution, likely altering the conformation dramatically.

**Table 1.**
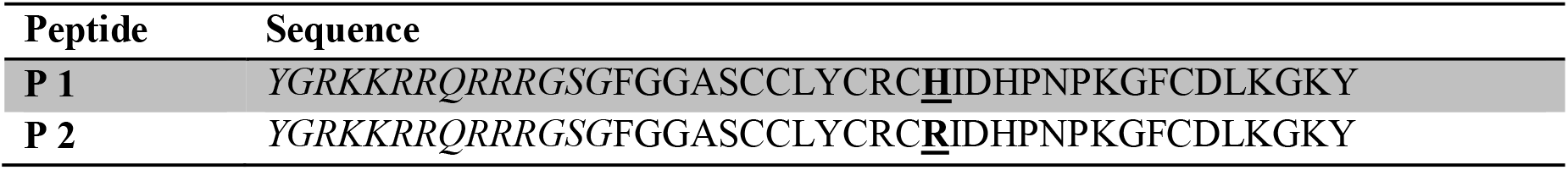
Amino acids sequences of Peptides P1 and P2 derived from the NSP10 protein of SARS-CoV-2 involved in the interaction with NSP16. Italics indicate amino acids of HIV-Tat sequence required for membrane penetration. Bold and underlined letters indicate substitution of Histidine with Arginine at position 26, corresponding to amino acid residue 80 in full length NSP10 (H80R).

**Figure 1.**
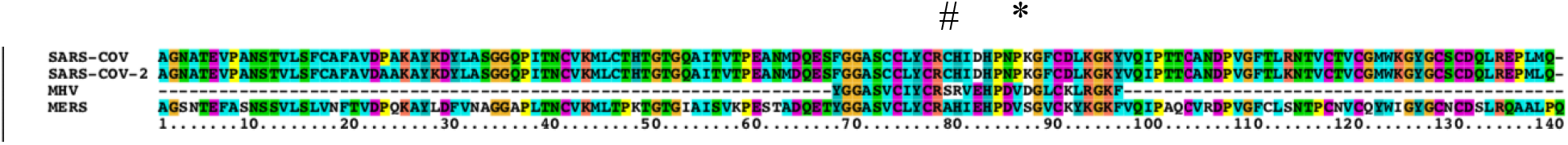
Alignment of the NSP10-derived sequence employed to inhibit the replication of Murine Hepatitis Virus (MHV) with full length NSP10 sequences from SARS-CoV, SARS-CoV-2, and MERS-CoV. * indicates a Proline to Valine substitution in MHV and MERS-CoV compared with SARS-CoV and SARS-CoV-2; # indicates residue His80 in SARS-CoV and SARS-CoV-2;

### SARS-CoV-2 NSP10-derived peptide (P1) induces secretion of IL-6 by human lung cells

We first determined the effect of peptides P1 and P2 on the secretion of IL-6 in human cell lines. As shown in Figure 2, incubation of the human NSCLC cell line NCI-H1792 with P1, but not P2 or a TAT sequence peptide only, resulted in a more than a 4-fold stimulation (P<0.001) of the intrinsic secretion of IL-6. Similar results were obtained with the human metastatic breast cancer cell line MDA-MB-231 (Figure 4)

**Figure 2.**
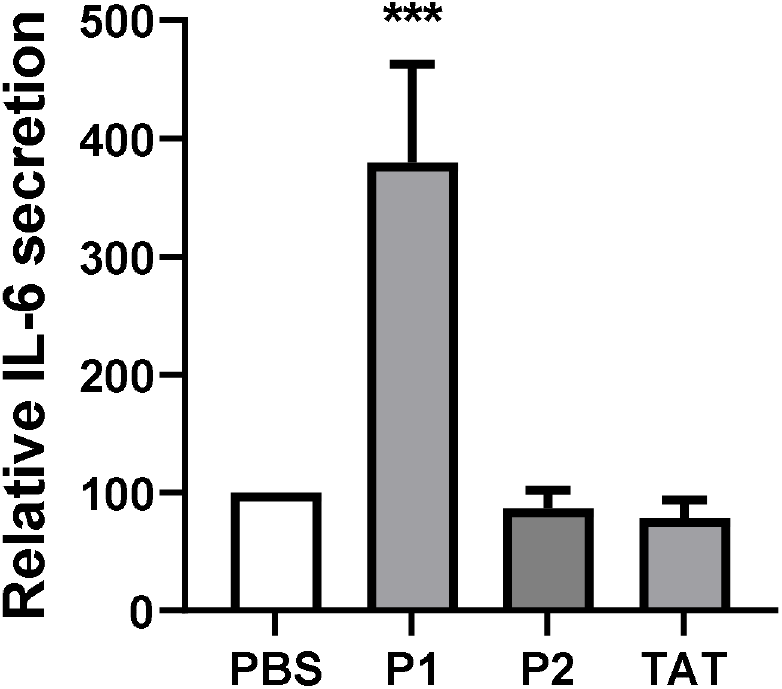
SARS-CoV-2 NSP10-derived sequences cause an increase in IL-6 secretion by human lung cancer cells. The human NSCLC cell line NCI-H1792 was incubated in the presence of NSP10-derived peptides P1 or P2, harbouring an N-terminal HIV-TAT sequence, or the TAT-only peptide for 24 h. The secretion of IL-6 was measured by ELISA and normalized to the control (PBS). Shown are the average results of 3 independent experiments. *** P<0.001.

### Plant-produced IL-37b and IL-38 effectively attenuate the levels of IL-6 induced by P1

We next determined the effect of plant-produced anti-inflammatory cytokines IL-37b and IL-38 on the P1-elicited stimulation of IL-6 secretion by H1792 cells. Pre-treatment with IL-38 resulted in pronounced attenuation (−60 %; P<0.001) of IL-6 secretion triggered by P1 (Figure 3). This attenuation effect, although less strong, was also observed when cells were pre-treated with IL-37b (−40%; P<0.05), or with a combination of IL-38 and IL-37b (−50%; P<0.05). Similar results were obtained using MDA-MB-231 cells. As shown in Figure 4, the attenuating effect of IL-38 and IL-37b was dose-dependent.

**Figure 3.**
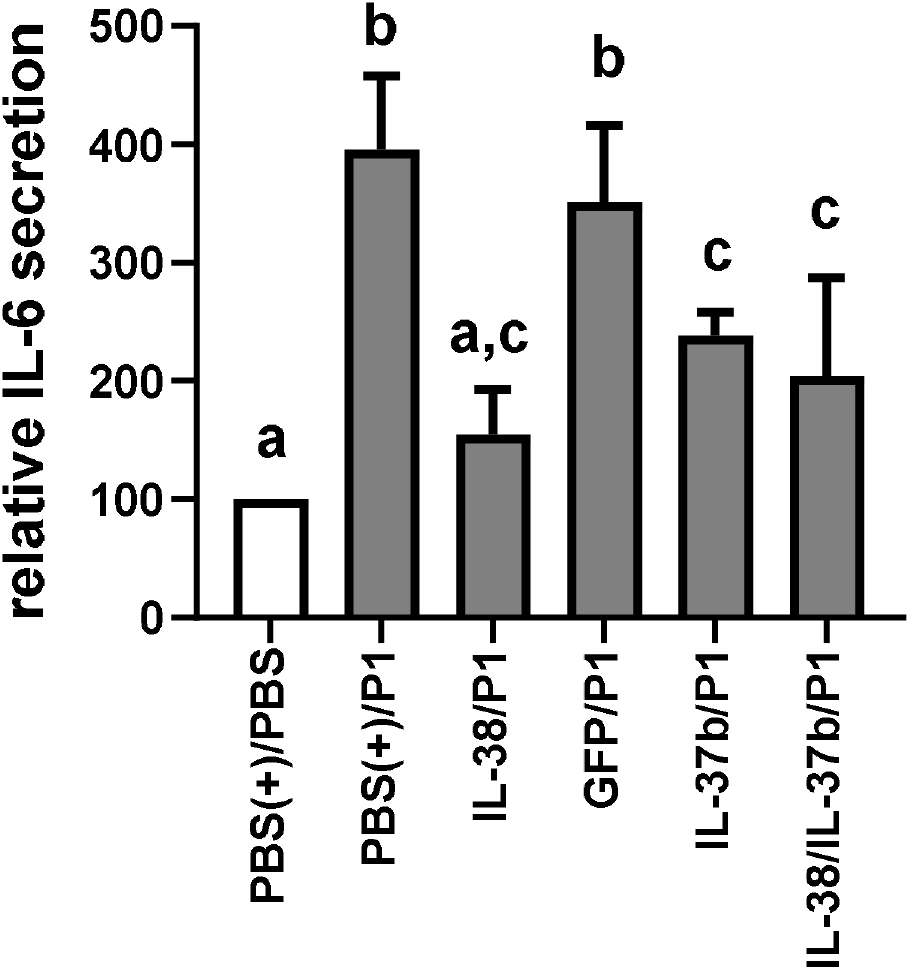
IL-37b and IL-38 attenuate the Peptide 1-induced stimulation of IL-6 secretion by human cells. IL-6 secretion by the human NSCLC cell line NCI-H1792 into the conditioned media was measured by ELISA. The cells were pre-incubated for 3 h in the presence of plant-produced recombinant IL-38, IL-37b (1.0 ng/μL), or a combination of IL-38 + IL-37b (0.5 ng/μL each), as indicated. PBS (1X) and a His-tag containing plant-produced GFP were used as controls. IL-6 secretion was stimulated by the addition of NSP10-derived Peptide 1 (10 μM) for 24 h. Shown are the average results of 4 independent experiments. Different letters above the columns indicate significant differences.

**Figure 4.**
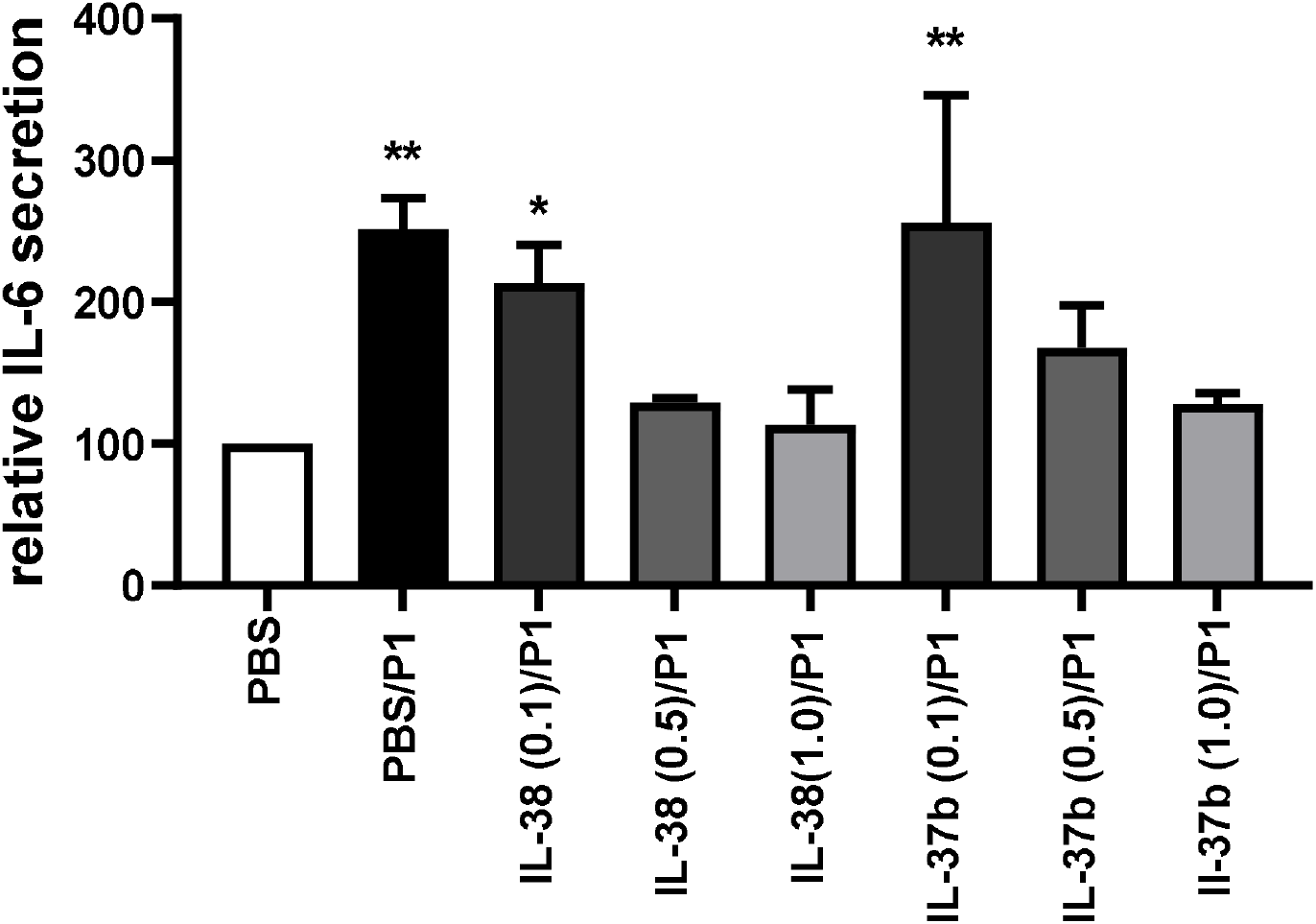
IL-37b and IL-38 attenuate the Peptide (P)1-induced stimulation of IL-6 secretion by human cells in a dose-dependent manner. IL-6 secretion by the human metastatic breast cancer cell line MDA-MB-231 into the conditioned media was measured by ELISA. The cells were pre-incubated for 3 h in the presence of plant-produced recombinant IL-38 or IL-37b at the indicated concentrations (ng/μL) prior to stimulation with peptide P1 (10 μM). PBS (1X) was added to the control. Shown are the average results of 3 independent experiments. ** P<0.01; * P<0.05, compared with the control.

### P1 and P2 cytotoxic activity

As the next step in the assessment of the biological action of peptides P1 and P2 we determined their cytotoxicity profiles, considering that NSP10 was shown to be implicated in multiple interactions with various host cell and viral proteins(Bouvet et al., 2014). Since Caco-2 cells are now routinely used in studies to examine potential drugs against SARS-CoV-2 due to viral preference for replication(Cagno, 2020), we tested the toxicity of P1 and P2 against Caco-2 cells. Our results showed that while P2 had no detectable toxicity at concentrations up to 200 μM, P1 displayed a Cytotoxic Concentration (CC_50_) of 11μM.

## DISCUSSION

### Designed peptides and cellular responses

The region of NSP10 corresponding to peptide P1 has previously been shown to inhibit complex formation between NSP10 and NSP16 from SARS-CoV(Ke et al., 2012). In another study, NSP10-derived peptides from MHV were used to demonstrate reduction in viral pathogenesis in cell lines and in mice(Wang et al., 2015). In the present study we have designed the P1 peptide based on the sequence of NSP10 from SARS-CoV-2 (identical in NSP10 from SARS-CoV) that is homologous to the MHV NSP10 sequence.

As a part of the NSP10 structure, the sequence of P1 contains a zinc knuckle motif consisting of a zinc ion coordinated by three cysteines and a histidine residue. Interestingly, only P1, but not P2, stimulated IL-6 production and caused cytotoxicity in our experiments. Peptide P2 differs from P1 in only one amino acid, replacing the histidine residue from the zinc finger motif by arginine. Our results therefore suggest that the zinc coordination site, which is greatly affected by the substitution of His by Arg in peptide P2, is needed for the induction of IL-6 expression. This could explain why P2 does not induce IL-6 in the same manner as P1. In a series of experiments to be reported elsewhere, we were in fact able to show that a peptide corresponding to a different interface region of the NSP10/NSP16 complex did not induce IL-6 and indeed reduced replication of SARS-CoV-2 in Caco-2 cells infected with the virus.

As seen in Supplemental Figure 2, the structure of MERS NSP10, which shares the Proline to Valine substitution with MHV, is slightly different from SARS-CoV and SARS-CoV-2 NSP10. While no experimental structure of the NSP10 protein of MHV exists, the MERS structure is available (PDB: 5YN5) as well as the SARS-CoV-2 NSP10 structure (DPDB ID:6W4H). We performed Normal Mode Analysis (NMA) using the DynaMut server on both structures to examine the areas of atomic fluctuations. The NMA results showed that MERS NSP10 has more deformation energies around the zinc knuckle, making the MERS NSP10 structure less stable around the Zinc finger (Supplemental Figure 3). Moreover, the residue corresponding to His80 in NSP10 from SARS-CoV, SARS-CoV-2, and MERS is an arginine in MHV. Taken together, this could provide an explanation as to why MHV NSP10 does not elicit an increase in IL-6 once applied to cells and mice. An alternative explanation could be that the reaction of murine cells is different from human cells. An experiment in which the levels of IL-6 are measured in mice and Caco-2 cells due to exposure to MHV NSP10-derived peptides, as well as SARS-CoV-2 NSP10-derived peptides would be beneficial to shed light on this inconsistency.

### Possible mechanism of IL-6 induction by the NSP10-derived P1

Host-virus interactome derived from proteomics and co-immunoprecipitation assays have suggested that NSP10 inhibits the NF-κB-repressing factor (NKRF) to facilitate interleukin-8 (IL-8) induction, and possibly IL-6(Li et al., 2021). This could potentially contribute to interleukin-mediated chemotaxis of neutrophils and the over-exuberant host inflammatory response observed in COVID-19 patients(Li et al., 2021). The link between NF-κB and IL-6 stimulation has been previously well-established. NF-κB is known to regulate an IL-6 mRNA stabilizing protein, known as AT-rich interactive domain-containing protein 5a (Arid5a)(Nyati et al., 2017). Toll-Like Receptor 4 (TLR4) induces NF-κB, which in turn activates Arid5a and subsequently induces IL-6 expression. While no previous studies examined whether NSP10 has a direct role in activating TLR4, the S protein of SARS-CoV-2 has been linked to TLR4(Brandão et al., 2020). There have been suggestions in the literature that SARS-CoV-2 non-structural proteins affect Toll-Like Receptors in general. In fact, both SARS-CoV and MERS-CoV viruses are known to affect TLR4 and TLR3(Totura et al., 2015). However, in the case of SARS-CoV and MERS-CoV, the mechanism appears to be protective against the viral infection.

### Curbing P1-induced IL-6 expression with plant-produced anti-inflammatory cytokines

Increased IL-6 expression in cells in response to P1 application could be curbed by treatment with recombinant, *in planta*-produced human cytokines IL-38 and IL-37b. It is reasonable to attribute the observed attenuation of IL-6 expression to the biological action of the cytokines, since treatment with the control protein GFP, bearing an identical His-tag, did not result in any significant attenuation. The anti-inflammatory action of both IL-38 and IL-37b was previously reported as dose-dependent, displaying bell-shaped concentration efficiency, with optimal concentrations ranging from 10 to 100 ng/mL(Nold-Petry et al., 2015; Van De Veerdonk et al., 2012). In addition, (Gu et al., 2015) showed that increasing concentrations of IL-37b (up to 500 ng/mL) applied in LPS-challenged THP-1 cells led to more significant inhibition of TNFα and IL-1β expression. In this regard, the pronounced attenuation of the levels of IL-6, an inflammation-associated marker, observed in our experiments is in accord with previous reports characterizing the anti-inflammatory nature of IL-38 and IL-37b at these concentrations (Cavalli and Dinarello, 2018; Gu et al., 2015; Han et al., 2020).

Despite their binding to a completely different set of receptors, the inhibitory action of both IL-37b and IL-38 was shown on the intracellular signal transduction pathways of different STATs, p38MAPK, ERK1/2 and JNK, as well as NF-κB signaling, outlining a degree of redundancy(Gao et al., 2021; Nold-Petry et al., 2015). IL-37 binds to II.-18Rα and recruits the IL-1R8 (also called SIGIRR or TIR8) to inhibit pro-inflammatory signalling(Nold-Petry et al., 2015). IL-37 also acts in the nucleus with Smad3 suppressing expression of inflammatory genes(Nold et al., 2010). IL-38 was first characterized as similar to IL-36Ra for its antagonist function on IL-36R, however, the mechanism of IL-38 signalling is not yet fully elucidated and appears to play a role in inflammation resolution(Van De Veerdonk et al., 2012). IL-38 has been shown to bind to three receptors: IL-36R, IL-1R1 and IL-1 receptor accessory protein-like1 (IL-1RAPL1), thus exerting anti-inflammatory effects by competing with their agonistic ligands and inhibiting their signalling pathways(Mora et al., 2016),(Yuan et al., 2016).

IL-38 produced a stronger inhibitory effect on IL-6 levels in comparison with either the action of IL-37b, or application of a combination of both recombinant cytokines (Figure 3). This observation is in agreement with their redundant inhibitory action on pro-inflammatory signalling leading to IL-6 expression, also indicating the absence of possible synergistic effects from the simultaneous application of IL-37b and IL-38.

### Conclusion

Our results demonstrated that a peptide derived from SARS-CoV-2 NSP10 can cause a significant increase of IL-6 secretion in human adenocarcinoma lung cells and resulted in cytotoxicity in an intestinal epithelial cell line. Our results also indicate that it is the zinc knuckle motif of NSP10, preserved in P1, but not in P2, that is likely to elicit this IL-6 inflammation marker increase response, emphasized by the notion that this motif is also found in other proteins known to cause increases in IL-6 expression. Since the elevated levels of IL-6 are a predictive molecular marker for the cytokine storm that accompanies COVID-19 pathology, correlating with poor prognosis, our findings suggest that therapeutic targeting the NSP10-induced inflammation should be pursued. IL-37b and IL-38 produced in engineered plant bioreactors were able to mitigate the induction of IL-6, suggesting that their application in the immunotherapy of COVID-19 should be further investigated.

## MATERIALS AND METODS

### Design of the peptides

The peptides were designed by using the homologous sequence of MHV’s NSP10 region interacting with NSP16 as this was the aim of inhibition. The HIV-Tat peptide sequence (YGRKKRRQRRRGSG) was added to the N-terminus. The peptides were modified with N-acetylation and C-amidation, artificially synthesized, purified using High Performance Liquid Chromatography (HPLC) and ensured not to have any disulfide bonds using Mass Spectrometry (MS) (Peptides 2.0 Inc). Prior to use, peptides were dissolved in 1XPBS.

### Toxicity tests of the peptides for Caco-2 cells infected with SARS-CoV-2 virus

Testing for toxicity of the peptides against cells infected with SARS-CoV-2 was done in a BSL-3 facility at Utah State University, part of the NIH/NIAID program. Confluent or near-confluent cell culture monolayers of Caco-2 cells were prepared in 96-well disposable microplates the day before testing. Cells were maintained in MEM supplemented with 5% FBS. The peptides were dissolved in 1XPBS and concentrations of 0.1, 1.0, 10, 100, and 200 μg/mL were prepared. Five microwells were used per dilution: three for infected cultures and two for uninfected toxicity cultures. On every plate controls for the experiment consisted of six microwells that were infected but not treated (virus controls) and six that were untreated and uninfected (cell controls). Peptide 1 and Peptide 2 were tested in parallel with a positive control drug using the same method as was applied for the peptides. The positive control was included with every test run. Growth media were removed and the peptides (0.1 mL) were applied to the wells at 2X concentration. Aliquots (0.1mL), containing virus at ~60 CCID50 (50% cell culture infectious dose) were added to the wells designated for virus infection. Media devoid of virus was added to the toxicity control wells and cell control wells. Plates were incubated at 37 °C with 5% CO_2_ until marked CPE (>80% CPE for most virus strains) was observed in virus control wells. The plates were then stained with 0.011% neutral red for two hours at 37°C in a 5% CO2 incubator. The neutral red medium was removed by complete aspiration, and the cells were rinsed 1X with PBS to remove residual dye. The PBS was completely removed, and the incorporated neutral red was eluted with 50% Sorensen’s citrate buffer/50% ethanol for at least 30 minutes. The dye content in each well was quantified using a microplate reader at 540 nm. The dye content in each set of wells was converted to a percentage of dye present in untreated control wells using a Microsoft Excel computer-based spreadsheet and normalized based on the virus control. The 50% effective (EC50, virus-inhibitory) concentrations and 50% cytotoxic (CC50, cell-inhibitory) concentrations were then calculated by regression analysis. It was not possible for us to compute the 50% effective (EC50, virus-inhibitory) since Peptide 1 was too toxic at 11μM while Peptide 2 had no detectable effect against the virus-infected cells even at 200 μM concentration, albeit being non-toxic at that concentration.

### Production and purification IL-37b and IL-38

Recombinant human cytokines IL-37b and IL-38 (UniProt identifiers Q9NZH6 and Q8WWZ1, respectively) were produced *in-planta* by Solar Grants Biotechnology Inc. using proprietary methodologies that will be discussed elsewhere due to pending patent applications. Briefly, the cytokines were expressed in their mature forms (V46 – D218 for IL-37b, C2–W152 for IL-38) with a C-terminal HIS-tag, purified using immobilized metal-affinity chromatography (IMAC, His SpinTrap Kit, GE Healthcare) and 0.22 μm-filtered (Millipore-Sigma) to obtain sterile solutions. The predicted molecular masses (20.3 kDa for IL-37b; 18.3 kDa for IL-38) were observed following Western blotting with protein-specific antibodies (MyBioSource Inc.). Native tetriary conformation was confirmed for each of the cytokines using protein-specific ELISA tests (R&D Systems Inc.).

### Cell culture for IL-6 measurements

The human non-small cell lung cancer cell line NCI-H1792 was obtained from ATCC. Cells were grown in RPMI Medium 1640 (Gibco) supplemented with L-glutamine, 10% fetal bovine serum (Gibco), and 1% penicillin/streptomycin at 37°C in a humidified atmosphere containing 5% CO2. For cell stimulation experiments, cells (5 x 104 per well) were seeded into 24-well plates and grown overnight. The next day, the cells were washed with OPTi-MEM (Gibco) and then pre-treated with plant-produced IL-38 (10 uL), IL-37b (10 uL), IL-38 plus IL-37b (5 uL each), or plant-produced GFP (1 ug) in 500 uL OPTi-MEM for 3 hrs at 37 oC. PBS (containing 25% glycerol) was added to the controls. Subsequently, peptides (10 uM) were added and incubation was continued for 24 h after which the conditioned media was harvested for determination of IL-6 secretion by ELISA.

### ELISA assays

Correct folding of plant-produced IL-38 and IL-37b anti-inflammatory cytokines was assessed by enzyme-linked immunosorbent assay (ELISA) using the DuoSet Human IL-38/IL-1F10 kit (R&D systems) and the Human IL-37/IL-1F7 uncoated ELISA kit (Invitrogen), respectively, following the instructions of the manufacturers. Secreted IL-6 was assessed using the Human IL-6 Uncoated ELISA kit (Invitrogen).

## ACKNOWLEDGEMENTS

This work was funded by the Federal Economic Development Agency for Southern Ontario (FedDev Ontario), through the Southern Ontario Network for Advanced Manufacturing Innovation (SONAMI), project number FC.2 *The computational work within this research was enabled in part by support provided by SHARCNET (http://sharcnet.ca) and Compute Canada (www.computecanada.ca)*.

