## Supplemental Tables and Figures for "Prolific Induction of IL-6 in Human Cells by SARS-CoV-2-derived Peptide is Attenuated by Recombinant Human Anti-inflammatory Cytokines made *in planta*"

Supplemental Information:

|  | Wt | H80R |
| --- | --- | --- |
| 0 | 0 | 0 |
| 1 | 0 | 0 |
| 2 | 0 | 0 |
| 3 | 0 | 0 |
| 4 | 0.195 | 0 |
| 5 | 1 | 0 |
| 6 | 1 | 0 |
| 7 | 1 | 0 |
| 8 | 1 | 0 |
| 9 | 1 | 0 |
| 10 | 1 | 0 |
| 11 | 1 | 0 |
| 12 | 1 | 0 |
| 13 | 1 | 0 |
| 14 | 1 | 0 |

Supplemental Figure 1: Occupancy of the Zinc ion in the Zinc knuckle motif with respect to the pH values. Data is derived from MultiConformer Continuum Electrostatics (MCCE) Monte-Carlo Simulation where the y-axis corresponds to occupancy from 0 to 1 and the x-axis corresponds to pH values. The H80R mutation causes a dramatic change in the electrostatic environment resulting in a large change in the occupancy.

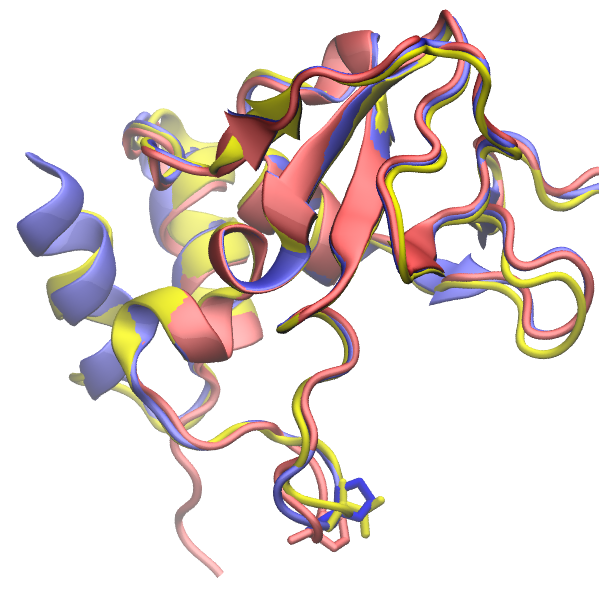

Supplemental Figure 2: The structure of nsp10 of SARS-CoV (blue) SARS-CoV-2 (magenta) and MERS-CoV (yellow). The mutation from Proline to Valine is shown in red circle.

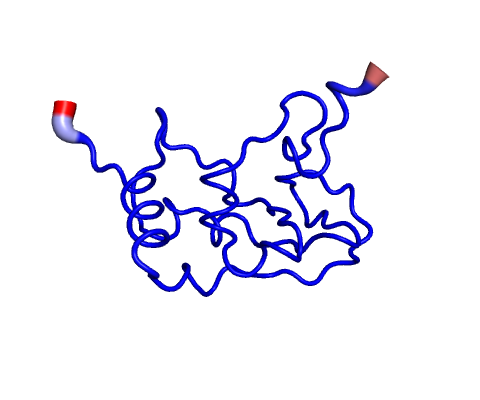

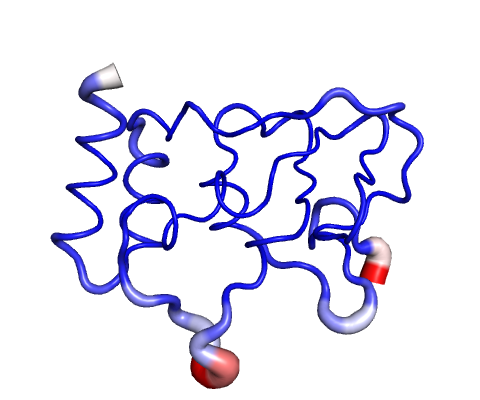

Supplementary Figure 3: Atomic fluctuations as measured by the DynaMut server. Left is MERS, right is SARS-CoV-2. Red indicate instability while blue indicates stability. Red circle indicates Zinc knuckle motif.
